# Data-driven Sampling Strategies for Fine-Tuning Bird Detection Models

**DOI:** 10.1101/2025.10.02.679964

**Authors:** Corentin Bernard, Ben McEwen, Benjamin Cretois, Hervé Glotin, Dan Stowell, Ricard Marxer

## Abstract

Passive Acoustic Monitoring has emerged as a promising tool for collecting ecological data, particularly in the context of bird population monitoring. Bird species can be automatically identified using pre-trained models, such as BirdNET. The performance of these models can be significantly improved through fine-tuning with annotated samples recorded in the specific acoustic conditions in which the microphones are deployed. However, PAM collects vast amounts of data, and annotating bird vocalizations requires specialized expetise. As a result, only a very small portion of the recordings can be effectively labeled. Selecting the most relevant samples to annotate in order to maximize performance in model fine-tuning remains a significant challenge. First, a regularization technique addresses the challenge of class imbalance during model fine-tuning. Next, a data-driven methodology is developed, introducing the *influence score*, which quantifies the impact of individual training samples on model performance to inform sampling strategies. A linear model is proposed to estimate the influence score for generalization to unseen data. Finally, several sampling strategies are compared, based on acoustic indices and predictions of the pre-trained model. Together, these contributions enable the identification of efficient annotation strategies to overcome the challenges of limited annotation resources in large-scale passive acoustic monitoring.

## 1 Introduction

Passive acoustic monitoring (PAM) provides an efficient method for large-scale biodiversity monitoring over time and space [1]. Birds are especially well-suited for PAM due to their role as ecosystem indicators and their detectability via acoustic recorders [2].

The growing availability of large-scale audio data [3] has led to the development of deep learning methods for the automatic detection and classification of bird vocalizations [4, 5]. Pretrained models such as BirdNET [6] are now available as open-source tools and are widely used for the identification of a broad range of bird species.

However, the accuracy and generalizability of these models are affected by multiple factors. First, model performance is strongly influenced by the recording process. This includes factors such as the type of recording equipment used [7] and the distance between the sound source and the recorder [8, 9]. In addition, the acoustic properties of the environment can alter the recordings, since different habitat types (such as forests, grasslands, wetlands) exhibit varying sound attenuation characteristics across frequencies [10, 11]. Environmental noise is another major source of variability. Background sounds such as wind, rain, and overlapping vocalizations from other species can mask target signals, leading to reduced detection accuracy [12]. Biological factors further impacts model generalizability, as species distributions vary across sites and seasons [13]. More importantly, some bird species exhibit geographic variation in their vocalizations, referred to as regional dialects [14, 15, 16], which can reduce the accuracy of models trained on data from other populations. Finally, pre-trained models are typically developed using single-species focal recordings with weak annotations (such as those available from Xeno-Canto [3]) which may limit their ability to generalize to the complex, multi-species soundscapes commonly encountered in PAM.

A common approach to address this challenge is to adapt existing models using in-situ data collected from the target environment [17]. This typically involves using a pre-trained model to compute embeddings, intermediate representations of the audio, and subsequently training a new classifier with annotated recordings from the specific sites of interest. Several models have been pre-trained specifically for bird sound recognition and have demonstrated the effectiveness of this transfer learning approach [18, 19, 20].

However, PAM generates massive volumes of raw audio data, yet only an extremely small fraction of these recordings can realistically be annotated. The process of labeling bird vocalizations is indeed both time-consuming and requires specialized expertise in ornithology. This limitation is further exacerbated when the goal is to monitor a large number of bird species, as the annotation budget must be divided across classes. In addition to data volume, bird vocalizations tend to be sparse, with a large proportion of recordings containing little or no target signal. Moreover, species occurrence is highly imbalanced, with a few common species frequently present while many others are rare or infrequent [21]. Consequently, efficiently selecting the most relevant samples for annotation in remains a major challenge.

### Scenario of the study

This work is part of the European project TABMON (Towards a Transnational Acoustic Biodiversity Monitoring Network), which aims to deploy a network of microphones and detect bird species across Spain, France, the Netherlands, and Norway, with the same material and protocol as in the *Sound of Norway* project [22]. As a consequence, the study focuses specifically on European bird species.

In this study, we consider a realistic scenario focused on the identification of 106 bird species, with a limited annotation budget allowing for the labeling of at most 500 audio samples, each 3 seconds in duration. The 106 bird species considered in this study are species that the pre-trained model has already been trained on, and for which we aim to improve detection performance under passive recording conditions at specific sites. To reflect the challenging conditions of PAM, the sampling strategies are evaluated using a dataset of soundscape recordings with multiple annotated bird vocalizations, complemented by a dataset of environmental noise to simulate low vocal activity. Under this setup, the annotated budget of 500 samples represents only 0.2% of the total data volume.

Our objective is to determine sampling strategies for model fine-tuning that lead to the highest class-wise mean Average Precision (cmAP). We selected cmAP as the primary evaluation metric because it reflects both the model’s ability to detect true positives (recall) and to avoid false positives (precision) across all classes. This metric does not require setting a confidence threshold. It computes the Average Precision independently for each class and then averages across all classes, assigning equal weight to each. This makes cmAP a particularly conservative (and therefore challenging) metric in the presence of high class imbalance. This performance metric reflects the need for the detection model to perform equally well across all species, not just the most common ones, which is essential for bird monitoring based on presence/absence data and for building reliable species occupancy models.

### Contributions

In the context of fine-tuning pre-trained models with limited annotated data relative to the number of species and under severe class imbalance, this paper makes the following contributions:

1. We demonstrate that a simple technique, L2-SP regularization, effectively improves finetuning for bird detection models. This technique mitigates catastrophic forgetting, a phenomenon in which a neural network loses previously learned knowledge while adapting to new data.
2. We develop a new data-driven methodology based on reverse correlation, introducing the *influence score*, which quantifies the impact of individual training samples on model performance. We used the influence scores to guide sampling strategies and to assess the performance ceiling given a fixed number of training samples. We also propose a linear model to estimate the influence score on unseen samples from their embeddings, enabling direct data sampling based on the predicted influence score.
3. We provide the first comparison between traditional sampling strategies based on the predictions of the pre-trained model BirdNET, including model confidence and uncertainty, and strategies based on acoustic indices, directly computable from the audio signal.

This work focuses on the detection of bird vocalization with BirdNET, but the methodology can be applied to other taxons and models. The scripts and data are open-source and publicly available on GitHub.

## 2 Dataset

To evaluate our approach in a controlled manner, yet under conditions similar to those of PAM, we used data from the WABAD dataset [23]. WABAD consists of 1-minute soundscape recordings containing multiple, strongly annotated bird vocalizations. As existing pre-trained models have not been trained on this dataset, it serves as a suitable and unbiased benchmark for this work. We selected only recordings from European sites. The files were split into train (40%), validation (10%) and test sets (50%). The recordings were cut into 3-second samples without overlap to match BirdNET’s input length. Annotations were attributed to segments when species vocalizations overlapped with the segment duration by at least 0.5 seconds. We retained only classes of bird species with at least one occurrence in each set. This led to a dataset of with ^13400 3^-sec samples in the train set, 3200 in the validation set and 16100 in the test set, with multi-labels corresponding to 106 bird species.

As presented in Figure 1.a, the class distribution in the dataset is highly imbalanced, with the the most common species appearing in over 2000 samples, while the rarest occurs in only a single sample.

**Figure 1.**
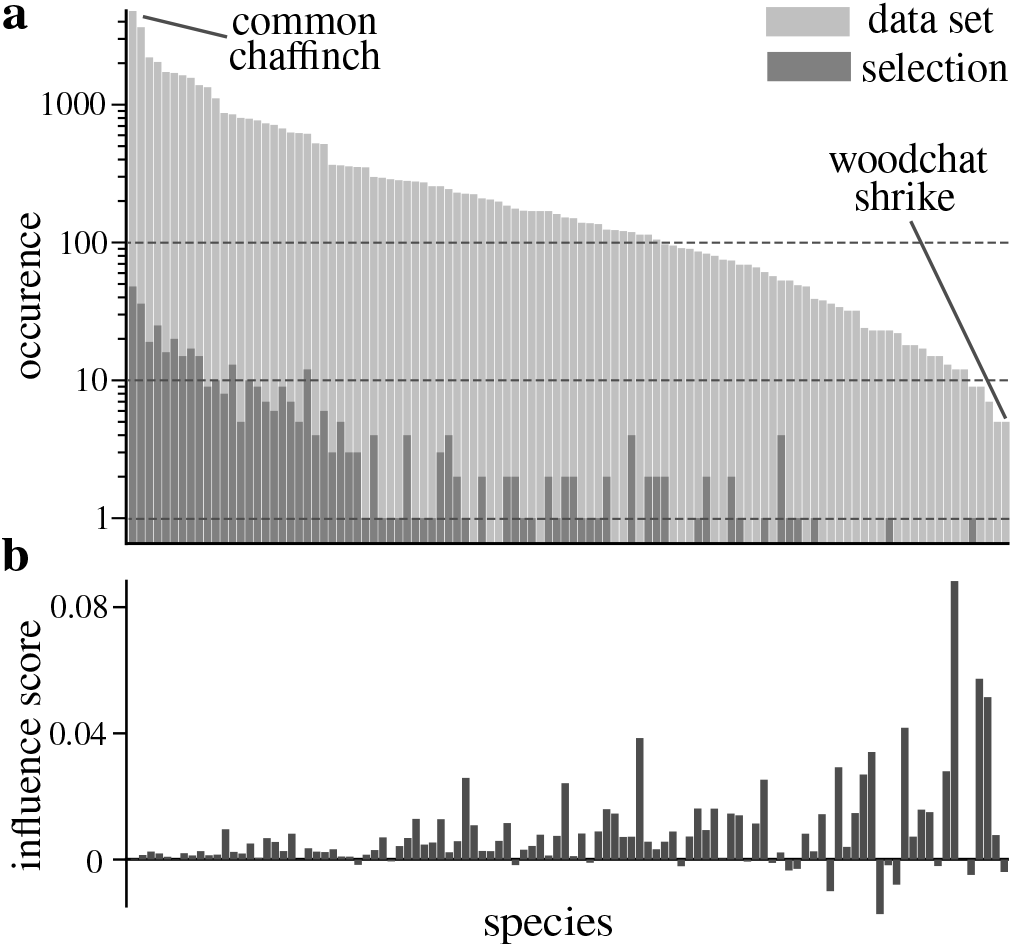
**a**. Species occurrence distribution showing high class imbalance. Each bar indicate the number of 3-second samples, in log-scale, of a given species in the full training set (light grey) and in a randomly selected fine-tuning set of 500 samples corresponding to the annotation budget (dark grey). Species are sorted on the x-axis according to their occurrence in the training set. Some species may be underrepresented or entirely absent from the fine-tuning set. **b**. Bars indicates the average influence scores of the species. Samples containing rare species show highest influence score.

In PAM, bird vocalization occurrences are often sparse, and only a small fraction of the recordings actually contain bird vocalizations. Some WABAD recordings contain sparse vocalizations, resulting in 3200 negative samples (i.e., segments taken between vocalizations) in the training set, accounting for just 20% of the samples. To simulate passive recording conditions, we incorporated noisy samples without bird vocalization into the dataset. Environmental noise was sourced from the ESC-50 dataset [24], including sounds produced by other animals (e.g., mammals and insects, with bird-related classes excluded), natural phenomena (e.g., wind, rain), and human-made sources (e.g., footsteps, engine noise). The recordings were segmented into 3-second chunks, with 2 sec overlap, resulting in the addition of 5500 negative samples to the 13400 samples of the training set. In the comparison of different sampling strategies (Sections 5 and 6), noisy samples from ESC-50 were further upsampled.

## 3 Model fine-tuning with highly imbalanced classes

Fine-tuning is commonly used in deep learning to adapt a pre-trained model to a more specific target task. This is usually done by training a new classification layer, with new data, on top of the existing embeddings. While this approach preserves the valuable intermediate representations learned by the pre-trained model, it forgets useful information related to species classification contained in the model’s original final layer. This problem, know as *Catastrophic forgetting*, typically occurs when a neural network learns new tasks and, in the process, forgets previously learned tasks. Several strategies have been proposed in the deep learning literature to mitigate this issue. These include architectural approaches that establish connections between the pre-trained and new model [19], data-based techniques that mix previously seen examples with new data [25], or regularization-based methods that add penalty terms to the loss function to preserve knowledge of the original task [26].

In bioacoustics research, fine-tuning pre-trained models, particularly BirdNET, is a widely adopted approach to improve the detection performance of poorly recognized species or to enable the detection of new ones. Despite this, strategies to address catastrophic forgetting are not yet standard practice. The process involves usually a limited number of species and requires a newly annotated dataset containing dozens to hundreds of examples per class [17]. In the case of large-scale PAM, which can involves a large number of species with high class imbalance and limited annotation resources, some species may be underrepresented or entirely absent from the fine-tuning set, as illustrated in Figure 1.a. This results in a degradation of the detection performances for these species with the fine-tuned model compared to the baseline performance of the pre-trained model.

This section optimizes a simple modification of the loss function, using L2-SP regularization, to counter catastrophic forgetting and improve the performance of bird detection models.

### Methods

Throughout this paper, BirdNET is used as pre-trained model, and model fine-tuning is performed by training a single linear classification layer on top of the BirdNET embeddings.

The L2-SP regularization [27] consist of incorporating a regularization term into the loss function ℒ to minimize the distance between the model weights **w**_*model*_ and the original weights of BirdNET’s last layer **w**_*birdnet*_:

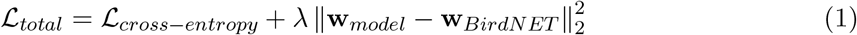

This prevents the model weights from deviating excessively from those of BirdNET. This method works for species that correspond to existing BirdNET outputs. The regularization parameter *λ* is a hyperparameter, optimized through grid search. We repeatedly selected *m* = 500 random samples from the training set and trained a model for *λ* values ranging from 10^*−*8^ to 10^2^, with 20 repetitions per condition, each using a different random seed for sample selection. Models were trained for a multi-label detection task using Adam optimizer and Binary Cross-Entropy loss, with a learning rate of 0.0001 over 150 epochs. We did not use a weighted loss function, commonly applied to imbalanced data, because it did not significantly improve performance in preliminary tests To accelerate convergence, the weights of the classification layer were initialized with those from BirdNET. Under these conditions, a single iteration of model fine-tuning took approximately 8 seconds when run on a GPU. After model training, the cmAP was computed on the validation set.

### Results

As illustrated in Figure 2, the results reveal the occurrence of *Catastrophic forgetting* under insufficient regularization strength. The optimal value for the regularization parameter was determined to be *λ* = 0.00032, and this value was adopted in all subsequent experiments.

**Figure 2.**
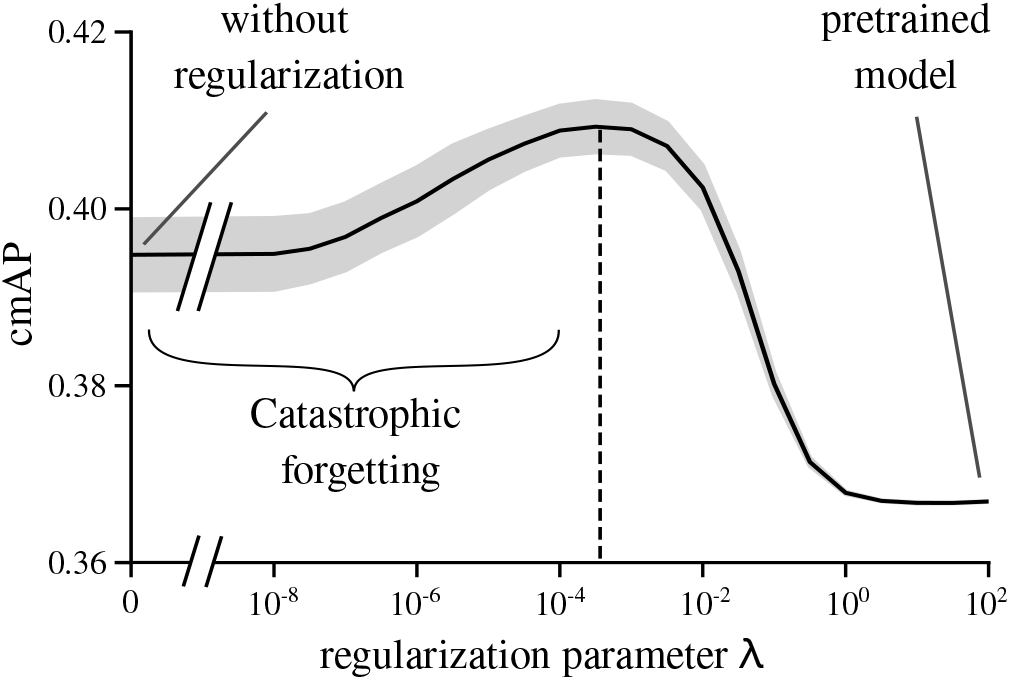
Effect of the regularization parameter *λ* on the class-wise mean Average Precision (cmAP) computed on the validation set. The black curve represents the mean, and the shaded area indicates the standard deviation across repetitions with different samples selection seeds. *λ* = 0 corresponds to no regularization, higher values of *λ* increasingly constrain the model to retain the pre-trained parameters. The maximum of the curve is used to determine the optimal value of this hyperparameter.

This method enabled us to improve the model detection performance and reach a cMAP=0.410 compared to a cMAP of 0.395 without regularization, and cmAP=0.367 for the pretrained model.

## 4 Reverse correlation for measuring sample influence scores

To improve model performance under a limited annotation budget, it is crucial to select training samples that provide the most informative value. In this section, we propose a new methodology based on reverse correlation [28], adapted from the field of model interpretability, to compute the *influence score* for each sample. This score quantifies the benefit of including a given sample in the fine-tuning set.

### Methods

This methodology consists of repeatedly performing the following procedure over a large number of iterations:

At each iteration *i*:

1. Samples selection: *m* samples are randomly selected from the training set to create a fine-tuning set, *m* corresponding to the annotation budget. A binary presence matrix *B* is constructed, where *b*_*i,j*_ = 1 if sample *j* is included in the selection at iteration *i* and *b*_*i,j*_ = 0 otherwise.
2. Model training: A model consisting of a single linear layer is trained on the fine-tuning set, using the training process and hyperparameters described in Section 3.
3. Model evaluation: The trained model is evaluated on the test set and the class-wise mean Average Precision (cmAP_*i*_) is computed as the performance metric.

After *N* iterations, we compute the influence score I_j_ of each sample *j* as the Pearson correlation between its presence vector **b**_*j*_ = [*b*_0,*j*_, *b*_1,*j*_, …, *b*_*N−*1,*j*_] and the vector of model performance scores **cmAP** = [cmAP_0_, cmAP_1_, …, cmAP_*N−*1_]. This measures the extent to which the inclusion of sample *j* in the fine-tuning set correlates with improved model performance.

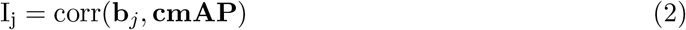

An informative sample is characterized by the model achieving slightly better performance when the sample is included in the fine-tuning set, and slightly worse performance than average when it is absent. As a result, its presence vector is positively correlated with the performance vector **cmAP**, leading to a high (positive) influence score. Conversely, a non-informative sample has little or even negative impact on model performance when included on the fine-tuning set, resulting in a low or negative influence score.

We applied this methodology on the 18900 samples in the training set (WABAD + ESC50), using an annotation budget of *m* = 500 samples per iteration and a total of *N* = 10000 iterations.

### Results

This methodology provided an influence scores for each sample. The influence score, averaged by species, is presented in Figure 1.b. The results show that samples containing rare species generally exhibit higher influence scores. However, this is not always the case, as some rare species are associated with negative influence scores, indicating that model performance is slightly below the average when these samples are included in the fine-tuning set. This may be due to a distribution shift between the training and test sets for these species, or to interactions arising when multiple species are present within the same sample.

Based on these results, we fine-tuned a model using the top 500 samples with the highest influence scores. This resulted in a detection performance of cmAP_*top*500_ = 0.489. This value serves as a topline for the best performance achievable when fine-tuning the BirdNET model with an annotation budget of *m* = 500 samples.

The influence score provides valuable insight into the potential benefit of annotating a given sample. However, the reverse correlation methodology is not directly applicable to new data in real-world PAM scenarios, as it requires fully annotated datasets. To assess the influence score for new data, we constructed a simple linear model to predict the influence score of a sample based on its BirdNET embedding. We used a ridge regression (implemented via scikitlearn), a linear regression method with L2 regularization that is effective for handling input with multiple variables. To avoid overfitting, the model was trained on half of the training set and then used to predict influence scores for the other half, called the sampling set. The L2 regularization strength was optimized via leave-one-out cross-validation and found to be *α* = 1800. The model achieved regression coefficients of *R*^2^ = 0.13 on train set (half used for training the ridge regression), and *R*^2^ = 0.08 when predicting the influence scores on the sampling set (the held-out validation half).

Although the regression coefficients were low, likely due to variability in the reverse correlation method, fine-tuning a detection model using the top 500 samples with the highest predicted influence scores from the sampling set resulted in a significant performance improvement, achieving achieving a cmAP = 0.410. For comparison, selecting the 500 samples with the highest influence score calculated with reverse correlation on the same sampling set led to cmAP = 0.452.

## 5 Data sampling based on acoustic indices

This section explores the potential of acoustic indices computed directly from raw audio signals without requiring prior annotation, to serve as effective sampling parameters Acoustic indices are widely recognized as effective tools for characterizing soundscapes, capturing aspects such as biodiversity and the relative contributions of anthropophony, biophony, and geophony [29, 30].

### Acoustic indices as proxy for the influence score

Results from the reverse correlation analysis were first used to assess which acoustic indices could be suitable candidates for data sampling by evaluating their potential as proxies for the influence score.

Acoustic indices were computed using the scikit-maad package [31]. For each 3-second audio sample in the dataset, spectral alpha indices were calculated using the *all spectral alpha indices()* function with parameters from [32]. Detailed descriptions of each acoustic index are available on the scikit-maad documentation page.

We then computed the absolute Pearson correlations between the acoustic indices and the influence scores I of the samples. The results are presented in Figure 3 (right). A high absolute correlation indicates that the corresponding acoustic index is an effective proxy for the influence score. We also grouped acoustic indices exhibiting similar behavior using agglomerative hierarchical clustering, as illustrated by the dendrogram in Figure 3 (left). The corelation distances between indices were defined as one minus the absolute correlation between them, computed across the dataset. Within each resulting cluster, we selected the index with the highest correlation with the influence score. This approach allowed us to retain a limited set of acoustic indices that are both mutually distinct and good proxies for the influence score.

**Figure 3.**
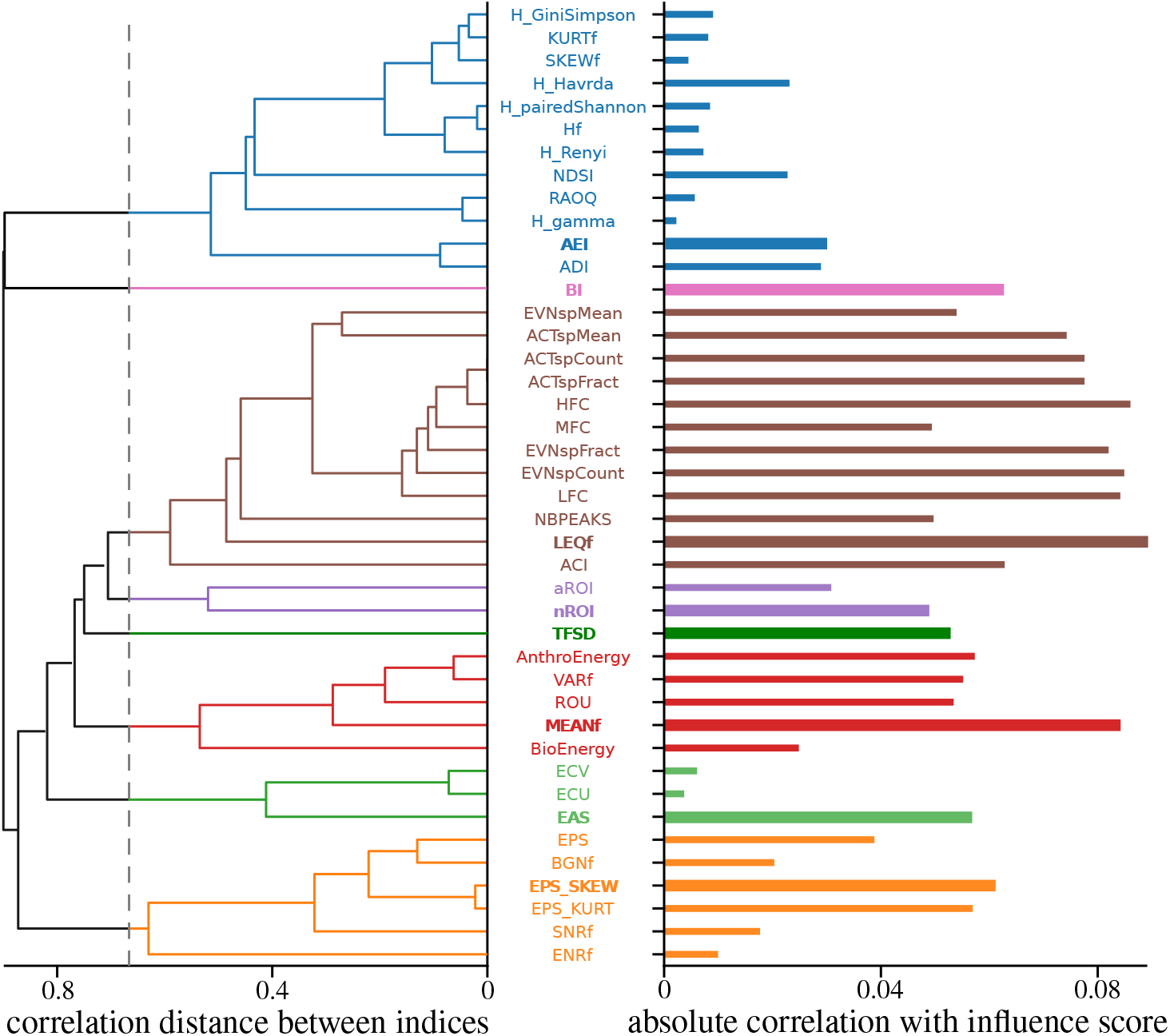
Left: Dendrogram showing the pairwise correlation distances between acoustic indices. The dotted line indicates the threshold chosen in order to cluster the indices into eights groups, each represented by different colors. Right: Bar plot showing the absolute correlation of each index with the influence score I. Within each cluster, the acoustic index with the highest absolute correlation is highlighted in bold. This enables the selection of acoustic indices that are distinct and reliable proxies for the influence score.

### Sampling Method

Following the identification of relevant acoustic indices, we assessed their effectiveness in guiding sample selection for annotation with PAM data. Specifically, we assessed threshold-based sampling strategies, in which instances are selected when the value of a given acoustic index exceeds (or falls below) a predefined threshold. Given the large number of available acoustic indices, we restricted the comparison to the subset previously identified as most relevant (in bold in Figure 3), and included the NDSI, a commonly used index, as an additional reference. For this experiment, we aimed to simulate more realistic bird call occurrences in PAM by targeting a scenario in which only 10% of the 3-second audio samples contains bird vocalizations. To achieve this distribution, samples from the ESC-50 dataset were upsampled such that environmental noise accounts for 90% of the training set, while the remaining 10% consists of bird vocalizations from WABAD. ESC-50 upsampling was achieved by assigning a higher drawing probability to noisy samples during the samples selection step.

For a given acoustic index value *α*_index_ and a threshold value *τ*_index_, the process was the following:

1. Samples selection: Samples are first selected from the training set based on whether their acoustic index satisfies the threshold condition *α*_index_ *> τ*_index_ (or *α*_index_ *< τ*_index_). Then, m = 500 samples are randomly drawn from this subset (with a higher probability for noisy samples). This creates the fine-tuning set.
2. *Samples annotation*: This step is here simulated by assigning the corresponding labels to the selected samples.
3. Model training: A model consisting of a single linear layer is trained on the fine-tuning set, using the training process and hyperparameters described in Section 3.
4. Model evaluation: The trained model is evaluated on the test set and the class-wise mean Average Precision (cmAP) is computed.

### Sampling Results

The results are presented in Figure 4. For each acoustic index, we can determine the most effective sampling strategy (*α*_index_ *> τ*_index_ or *α*_index_ *< τ*_index_) and estimate the threshold value that maximizes model performance. All the tested acoustic indices resulted in improved performance compared to random sampling. Selecting samples with a TFSD *>* 0.54 emerges as the most effective strategy for data sampling. TFSD (Time frequency derivation index) captures temporal and spectral modulations in the 2–10 kHz frequency range, which are indicative of bird vocal activity in the spectrogram. LEQf, MEANf, and EPS-SKEW, also show promising potential for guiding sample selection. The results obtained with LEQf (Equivalent Continuous Sound Level) and other sound level–based indices should be interpreted with caution, as they may reflect biases in signal intensity caused by differences in recording equipment and protocols across datasets.

**Figure 4.**
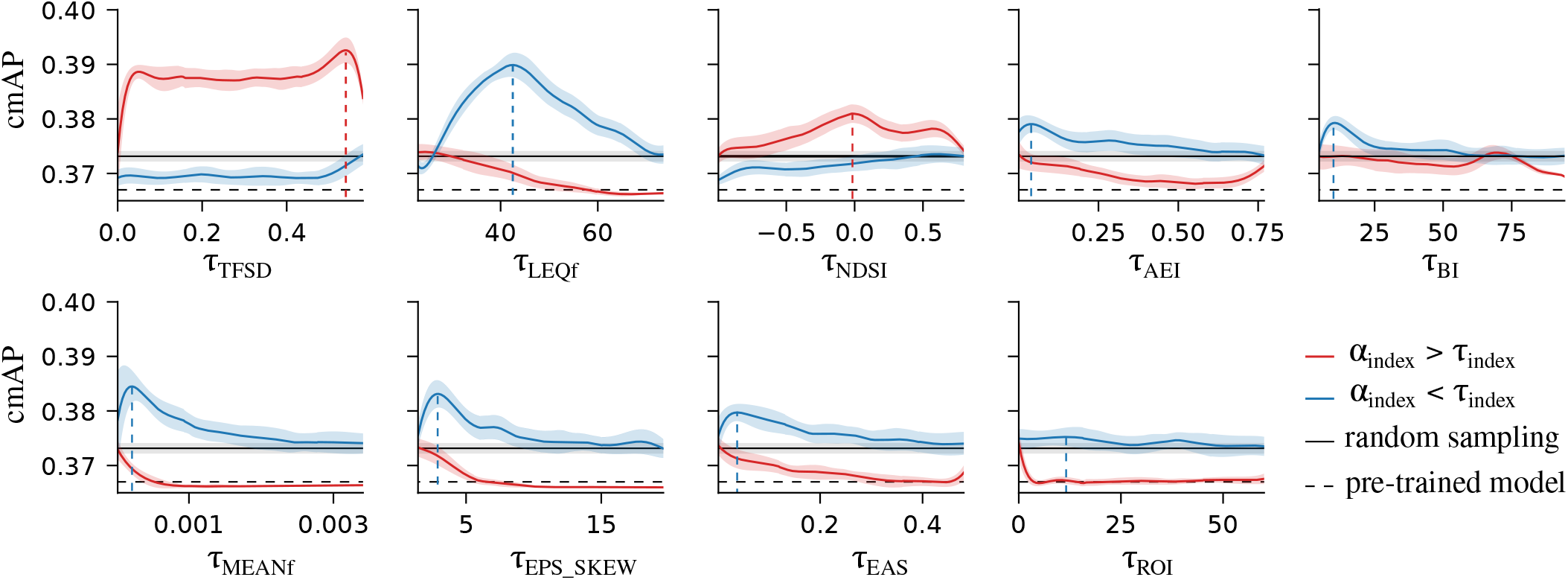
Evolution of the class-mean Average Precision (cmAP) for models trained using different dataset sampling strategies based on the values of the previously selected acoustic indices *α* and varying threshold values *τ*. Colored curves represent the mean cmAP across repetitions with different random sampling seeds, while shaded areas shows the standard deviation. The peak of each curve is used to identify the optimal threshold. As baselines, the solid black line shows the cmAP achieved with completely random sampling, and the black dotted line corresponds to the performance of the pre-trained BirdNET model.

## 6 Data sampling based on pre-trained model predictions

When the pre-trained model provides not only embeddings but also reasonably accurate predictions for the target classes, such as BirdNET for bird species present in the WABAD dataset, its is possible to leverage the predictions to guide sample selection.

### Sampling Method

We compared sample selection using two parameters derived from the confidence scores *p*_*k*_ output by BirdNET for each species *k*:

- The maximum confidence score across species: P_*max*_ = max_*k∈*[0:*K−*1]_(*p*_*k*_) This value reflects the presence of birds in a given sample, as higher scores typically indicate stronger evidence for at least one species.
- The maximum binary entropy across species:

H_*max*_ = max_*k∈*[0:*K−*1]_(*h*_*k*_) with *h*_*k*_ the binary entropy for species *k*: h_*k*_ = *−p*_*k*_ log *p*_*k*_ *−*(1*− p*_*k*_) log(1 *− p*_*k*_) This metric is commonly used in active learning to captures the model’s uncertainty [33]. A sample with high entropy indicates greater uncertainty from the model in making a prediction.

The evaluation process was exactly the same as in Section 5, and the samples were selected when the value of the parameter P_max_ or H_max_ exceeds (or falls below) a predefined threshold.

### Sampling Results

The results are presented in Figure 5. Selecting samples with a high maximal binary entropy (H_max_ *>* 0.65), indicating high model uncertainty for at least one class, emerges as the most effective sampling strategy. Interestingly, selecting samples with a maximum confidence score above a very low threshold (e.g., P_max_ *>* 0.08) leads to nearly comparable performance. This suggests that such a threshold is sufficient to capture bird vocalizations, in line with recent results [34]. In contrast, restricting selection to high-confidence samples appears to negatively impact model performance.

**Figure 5.**
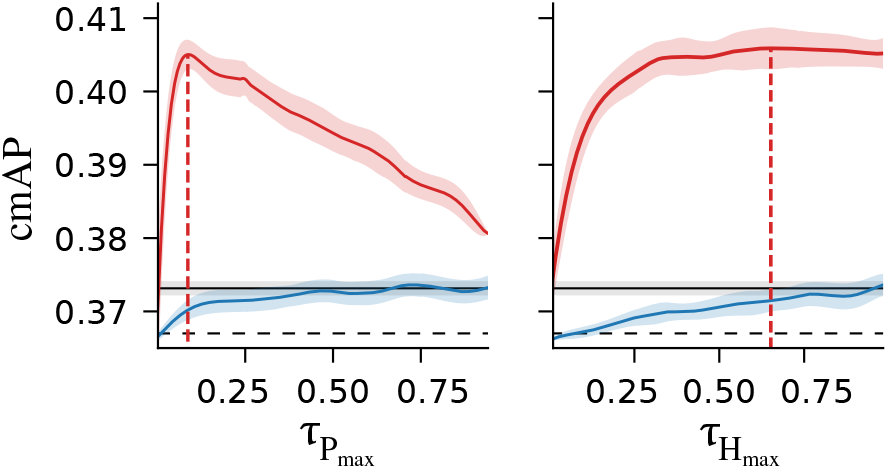
Evolution of the class-mean Average Precision (cmAP) for models trained using different dataset sampling strategies based the predictions of the pre-trained model and varying threshold values *τ*. The same color coding as in Figure 4 is applied.

## 7 Discussion

The best sampling strategies are summarized in Figure 6. Based on the predictions of the pre-trained model, maximal binary entropy, reflecting model uncertainty, appears as the most effective overall sampling strategy. Uncertainty sampling is a commonly used strategy in active learning [35, 36, 37], which is based on an iterative loop of sample selection and model training. While these approaches are effective when using small batch sizes, they require frequent interactions between the model and human annotators, something that is not always feasible in bird sound annotation, where expert knowledge is rare and the process can be time-consuming. Some acoustic indices also stand out as valuable for data sampling, such as TFSD. While acoustic indices yield lower performance than pre-trained model predictions, they remain valuable sampling parameters, as they can be directly computed from the raw acoustic signal. It is especially relevant in cases where the pre-trained model does not produce predictions, such as with unsupervised models [20] or when attempting to detect new species not covered by the pre-trained model.

**Figure 6.**
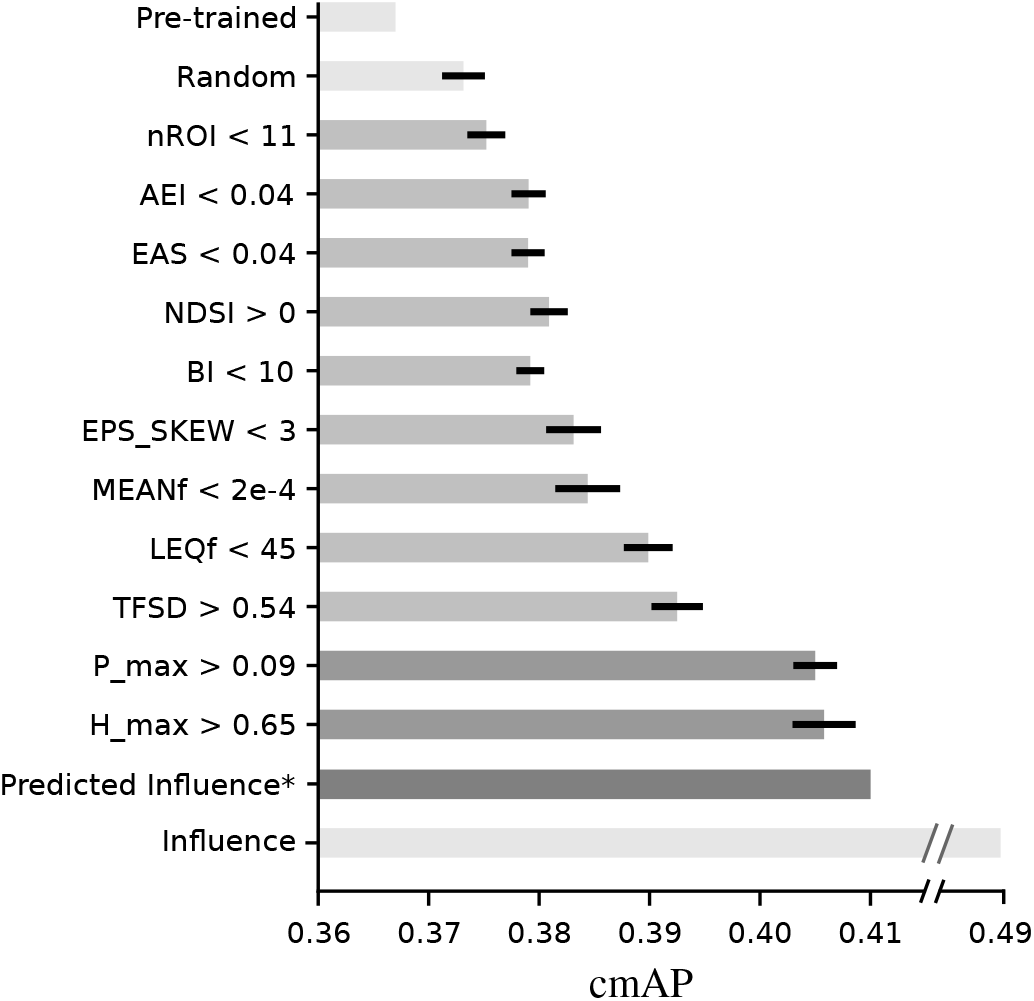
Summary of performances (cmAP) achieved by models trained on 500 samples selected using the best strategies. Error bars indicate the standard deviation across repetitions of the random sub-selection process using different seeds. The baselines are shown in light gray. The baselines correspond to the performances of the pre-trained BirdNET model without fine-tuning and a model trained on 500 randomly selected samples. The top is achieved by a model trained on the 500 samples with the highest influence scores. *Note: due to overfitting concerns, the strategy of selecting the 500 samples with the highest predicted influence scores was applied to only half of the training set, which likely underestimates the performance in comparison.

Although the overall cmAP gains may appear modest, they represent a valuable improvement. Average Precision, calculated here across 106 classes, is indeed a particularly conservative and challenging metric, especially in the presence of severe class imbalance.

In most strategies, we observe that the performance reaches a maximum at a specific threshold value, beyond which it declines as the threshold becomes more restrictive. This decrease is likely due to a reduction in data diversity. Effective model training typically requires a balance between informativeness and diversity of the training samples. In our current approach, diversity is maintained by randomly sampling from the subset that meets the selection criteria, introducing variability and reducing overfitting to homogeneous data. In future work, we plan to explore more advanced diversification methods [38] to further close the gap with the upper baseline, achieved by training on the samples with the highest influence scores.

The study presents however some limitations. First, the results may be biased by the characteristics of the dataset. Although the WABAD dataset includes recordings from 26 different sites across various European countries, this coverage may still be insufficient to fully capture regional variability and ensure generalizability. The recordings in WABAD were manually selected for annotation, which may introduce biases, such as the over-representation of certain species of interest. To simulate low bird call occurrence, we added and upsampled environmental noise in the training data using the ESC-50 dataset. This dataset contains approximately 40 types of noise, each represented by an equal number of examples. However, this noise distribution is unlikely to reflect real-world conditions in PAM. In practice, passive recordings from natural areas typically contain predominantly natural sounds, such as wind, rain, or extended periods of low acoustic activity, while human-made sounds are often rare. The overrepresentation of certain human sounds in ESC-50, such as laughing, clock alarms, or sirens, may significantly affect the performance of acoustic indices that are sensitive to the biophony (i.e., sounds in the 2–10 kHz range). Indices such as TFSD, NDSI, and BI could potentially perform better under more realistic conditions where natural ambient noise dominates.

Furthermore, acoustic indices are typically computed over longer time periods, often using recordings of one minute or more [39]. In this study, however, the indices are calculated at the sample level, using 3-second audio segments to align with the input length required by BirdNET. As a result, the ranges of temporal averaging in some indices are shorter than usual, introducing greater variability. Additionally, the recordings in the WABAD and ESC-50 datasets were captured using different recording equipment, with different gains and at varying sampling frequencies, which may further contribute to the variability of the acoustic indices.

## 8 Conclusion

In this study, we first demonstrated the benefit of incorporating a regularization term into the loss function to partially constrain the layer weights toward those of the pre-trained model. This approach led to significant improvements in model fine-tuning under the challenging conditions of large number of species, severe class imbalance, and limited annotations.

Based on reverse correlation, we developed a methodology to estimate the individual influence score of each training sample on model performance. These influence scores were then used to compute a topline, representing the maximum performance achievable by fine-tuning the model with the most influential samples. A linear model was trained to predict influence scores on unseen data from their embeddings, and the strategy of selecting samples with the highest predicted influence scores led to strong performance. Additionally, the results informed the selection of a limited set of distinct acoustic as strong sampling parameters candidates.

Finally, we compared different sampling strategies consisting on selecting samples above (or below) a given threshold, using acoustic indices or prediction of the pre-trained model. The best performance gain was obtained when selecting samples with a high model uncertainty, in line with previous literature on active learning. When the pre-trained model predictions are not available, such as with unsupervised models or for new species, the acoustic index TFSD appears as a promising sampling parameter that can be calculated directly on the acoustic signal.

In summary, we introduced key improvements to model fine-tuning and annotation sampling strategies, addressing the challenges posed by limited annotation resources and the presence of numerous, imbalanced species. Recommendations for sampling strategies according to the type of pre-trained model are summarized in Table 1. These contributions pave the way for more efficient models capable of analyzing recordings with spatial and temporal variations necessary for large scale passive acoustic monitoring networks for biodiversity assessment. Together, these contributions pave the way for more efficient models able to handle spatial and temporal variability in recordings, an essential requirement for large-scale passive acoustic monitoring networks in biodiversity assessment.

**Table 1:** Recommendations for annotation sampling strategies based on the model type.

| Pre-trained Model Conditions | Recommended Sampling Strategy |
| --- | --- |
| Pre-trained model that does not provide outputs (unsupervised). Pre-trained model that has not been trained on your target species. | Acoustic indices, such as TFSD above a threshold. (Note: strategies based on embeddings clustering may also be effective.) |
| Pre-trained model that provides predictions for your target species. | Model predictions, focusing on samples with high uncertainty or samples above a low confidence score. |
| BirdNET, with target species and distribution similar to those in this study | Predicted influence score, which can be computed using the provided linear regression model. |

## Acknowledgments

This research was funded by Biodiversa+, the European Biodiversity Partnership, in the context of the Towards a Transnational Acoustic Biodiversity MOnitoring Network (TABMON) project under the 2022-2023 BiodivMon joint call. It was co-funded by the European Commission (GA ref. 101052342) and the following funding organisations: Norwegian Research Council (project number 350977), l’Agence Nationale de la Recherche (ANR-23-EBIP-0010), and l’Office français de la biodiversité (OFB-23-1865). H. Glotin research is partly granted by ANR-21-CE04-0019 SYLVANIA.

## Author Declarations

### Data availability

The scripts and data are open-source and publicly available on GitHub.

